# LI Detector: a framework for sensitive colony-based screens regardless of the distribution of fitness effects

**DOI:** 10.1101/2020.06.27.175216

**Authors:** Saurin Bipin Parikh, Nelson Castilho Coelho, Anne-Ruxandra Carvunis

## Abstract

Microbial growth characteristics have long been used to investigate fundamental questions of biology. Colony-based high-throughput screens enable parallel fitness estimation of thousands of individual strains using colony growth as a proxy for fitness. However, fitness estimation is complicated by spatial biases affecting colony growth, including uneven nutrient distribution, agar surface irregularities, and batch effects. Analytical methods that have been developed to correct for these spatial biases rely on the following assumptions: i) that fitness effects are normally distributed, and ii) that most genetic perturbations lead to minor changes in fitness. Although reasonable for many applications, these assumptions are not always warranted and can limit the ability to detect small fitness effects. Beneficial fitness effects, in particular, are notoriously difficult to detect under these assumptions. Here, we developed the linear interpolation-based detector (LI Detector) framework to enable sensitive colony-based screening without making prior assumptions about the underlying distribution of fitness effects. The LI Detector uses a grid of reference colonies to assign a relative fitness value to every colony on the plate. We show that the LI Detector is effective in correcting for spatial biases and equally sensitive towards increase and decrease in fitness. LI Detector offers a tunable system that allows the user to identify small fitness effects with unprecedented sensitivity and specificity. LI Detector can be utilized to develop and refine gene-gene and gene-environment interaction networks of colony-forming organisms, including yeast, by increasing the range of fitness effects that can be reliably detected.

## INTRODUCTION

Colony-based high-throughput screens (CBHTS) of microbes are increasingly used for basic science biomedical and industrial research^1-4^. These screens involve growing manually or robotically “pinned” grids of microbial colonies on agar plates and recording colony growth using imagery. The images are computationally analyzed to generate a quantitative output of colony size, which is used as a proxy for the organism’s fitness. The wide availability of tools to conduct and analyze CBHTS, combined with the growing number of artificial gene constructs for microbial model organisms, has provided a large-scale controlled approach to experimentally determine the effects of genetic and environmental perturbations on the fitness of an organism. CBHTS have been used to explore genetic interactions^5,6^, protein-protein interactions^7-9^, chemical-genetic interactions^10-12^, and microbial pathogenicity^13^.

CBHTS fast track discovery thanks to the scale at which they are performed. However, spatial biases like edge effects^14,15^, local competition^15,16^, batch effects^15,17^, source plate memory^14^, light artifacts^18,19^, agar surface nutrient heterogeneity^14-16^ and humidity^20^, all lead to undesired colony size differences that are not relevant to the biological question being investigated (**Supplementary Fig. S1**). These spatial biases need to be corrected before making any biological inferences. The extent of spatial bias is difficult to predict *a priori*, making its identification and correction a substantial computational challenge^17^. A variety of existing tools implement normalization algorithms to correct for spatial biases, including the HT colony grid analyzer^21^, Colonyzer^18^, ScreenMill^19^, ScanLag^22^, SGATools^15^, Balony^16^, Scan-o-matic^23^, and MATLAB Colony Analyzer Toolkit (MCAT)^14^ (**Supplementary Table S1**).

Most of the existing tools rely on the following assumptions about the distribution of fitness effects (DFE): that the colony sizes in an experiment are normally distributed and that genetic manipulations rarely cause significant fitness deviation from wildtype^15^. These assumptions can be violated in experiments with biased sets of mutants^17,24^, or when experimental conditions produce a high variance in the DFE^17^. Even in unbiased genome-scale screens, the assumption of normal distribution is usually violated due to a skew towards negative fitness effects^25-27^. As a result, while existing methods can reliably detect large changes in fitness, they are less sensitive in detecting small effects which are difficult to differentiate from noise. This difficulty in detecting small fitness effects is especially pronounced for small increases in fitness. Overall, methods that rely on strict assumptions about the underlying DFE reduce the power of CBHTSs for broader scientific inquiry.

Here, we present the linear interpolation-based detector (LI Detector or LID), a CBHTS framework designed to avoid making any *a priori* assumptions about the underlying DFE. This two-part, experimental, and analytical framework utilizes a reference colony grid on every plate of the experiment to predict and correct for spatial biases. The reference grid is an isogenic population of colonies that are evenly distributed over the agar surface to act as internal local controls^23,28^. Our results show that the LI Detector’s reference colony based linear interpolant can successfully control for spatial bias. LI Detector is a tunable system that can provide the users with the ability to identify 5% or lower fitness effects with very high specificity and sensitivity. LI Detector performs as well as a popular existing method (MCAT^14^) when the underlying DFE is normal, and better when that is not the case.

## RESULTS

### Development of a new CBHTS framework

The LI Detector framework is specifically designed to correct spatial bias and sensitively detect small but significant fitness changes without making *a priori* assumptions about the underlying DFE of tested strains (thereafter, “mutant” strains). The experimental pipeline (**Fig. 1A**) follows a pin-copy-upscale protocol that serves two purposes. It reduces colony size differences that arise during the pinning process and adds a reference colony grid^23^ on every plate. The analytical pipeline utilizes the reference colony grid to correct spatial bias and infer the fitness of mutant strains relative to the reference strain. The analytical pipeline (**Fig. 1B**) consists of five main steps: 1) local artifact correction, 2) source normalization, 3) reference strain based background colony size estimation using a 2-dimensional linear interpolant, 4) estimation of spatially-corrected relative fitness as the ratio of the local artifact corrected colony sizes divided by the estimated background colony sizes, and 5) empirical hypothesis testing to identify mutant strains with colony size distributions that have a significantly larger or smaller mean than the reference strain.

**Figure 1.**
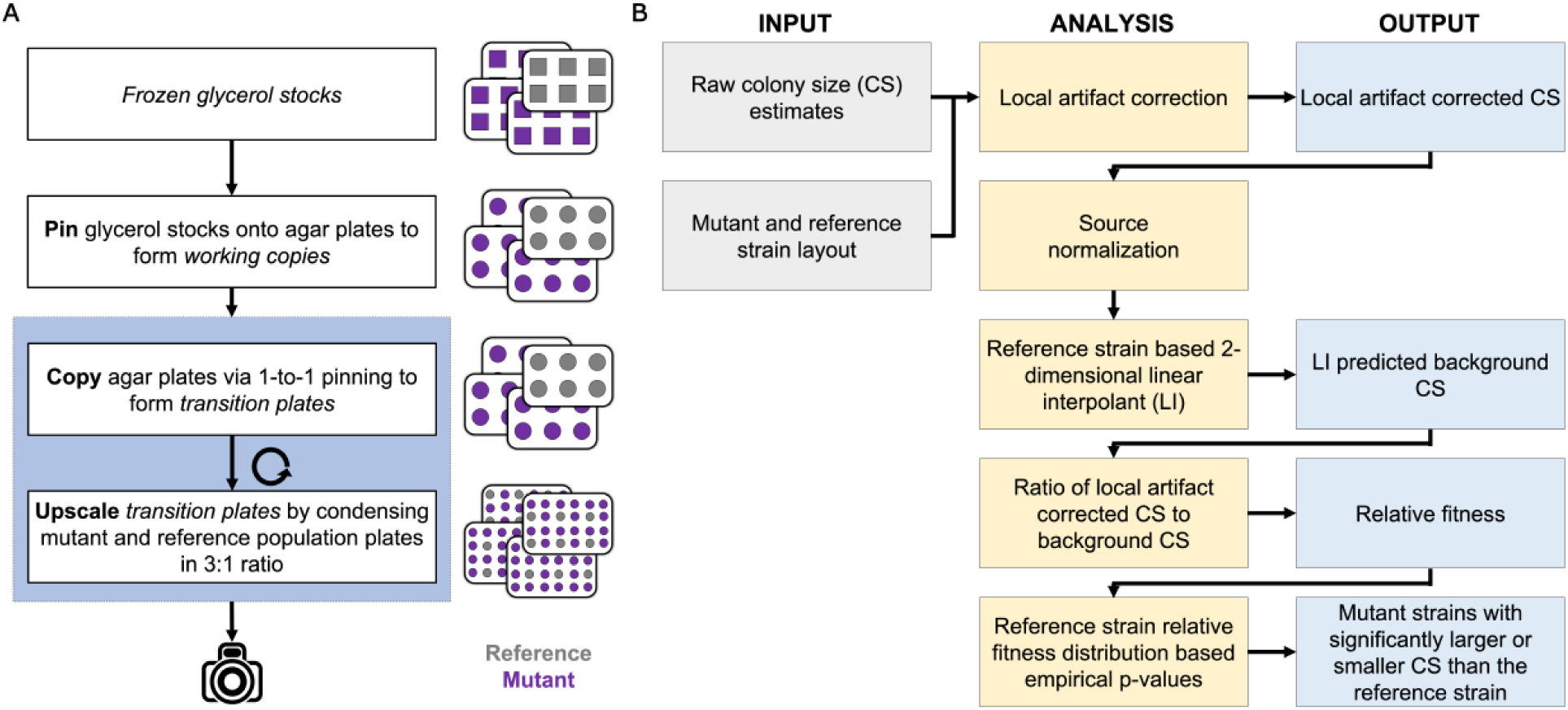
The LI Detector framework consists of integrated experimental and analytical pipelines. **A**. The pin-copy-upscale experimental pipeline from frozen glycerol stocks (top) to imaging (bottom). Each box represents a pinning step, and the steps within the sky-blue highlighted portion can be repeated until the desired colony density is reached. Illustrations to the right of the flowchart is a simplified representation of four experimental plates. A reference population (grey) is introduced on every plate during the first upscale step. The analytical pipeline uses this population for spatial bias correction and relative fitness estimations for the mutant strains of interest (purple). **B**. Workflow of the analysis pipeline where columns from left to right represent user inputs, analytical steps, and outputs. User inputs consist of raw colony size estimates and the strain layout of the plates. The analytical pipeline performs: i) local artifact correction, ii) source normalization, iii) reference-based background colony size estimation using a 2-dimensional linear interpolation, iv) corrects for spatial bias by dividing the local artifact corrected colony sizes with the background colony sizes and provides a measure of relative fitness, and iv) assigns empirical p-values using the reference strain relative fitness distribution. The outputs include local artifact corrected colony sizes, background colony sizes, spatially corrected relative fitness, and mutant strains identified as having a mean colony size that is significantly larger or smaller than the reference strain.

The local artifact correction step is designed to reduce spatially localized colony size differences that arise due to differential access to nutrients. It is similar to the competition correction feature implemented by several existing methods^15,16^. The source normalization step controls for differences in colony sizes that occur as a result of the upscaling process. This step was reimplemented from the interleaving feature of MCAT^14^. Briefly, it deconstructs the colony size estimates of the higher density plates into subsets corresponding to the source plates used for the upscaling (**Supplementary Fig. S2**). Both local artifact correction and source normalization are provided as optional steps in the LI Detector analytical pipeline.

The background colony size estimation step predicts what the size that a reference colony would be for every position of the plate. This step employs a 2-dimensional linear interpolant based on the reference colony grid. Relative fitness is then assigned to every colony as a ratio of the local artifact corrected colony size to the predicted background colony size. This estimate of relative fitness corrects local spatial bias without making any assumptions of the underlying DFE. The only assumption is that, for any location on the plate, the spatial bias is expected to affect the reference and mutant colonies to an equal extent.

Each mutant strain is assigned a relative fitness value corresponding to the average relative fitness of its replicate colonies. The distribution of relative fitness estimates for the reference strain is then used as null distribution to calculate empirical p-values describing the probability of the reference strain having a more extreme value of relative fitness than the mutant strain. The empirical p-values are used to determine the significance of the mutant strain fitness deviation from the reference strain (**see Materials and Methods**).

In what follows, we compare the performance of LI Detector with one of the most versatile and robust tools available for correcting spatial bias, MCAT^14^. The overall workflow adopted for testing the two methods’ performance is described in **Table 1** and **Supplementary Figure S3**. In brief, we estimated the specificity and sensitivity of the LI Detector and MCAT using colony size datasets generated using an isogenic population of *S. cerevisiae* (**see Materials and Methods**). A subset of colonies was mocked as references, and the rest were mocked as mutants. The LI Detector and MCAT spatial bias correction was applied independently. For consistency, our empirical p-value calculation strategy was used for the two methods. The mutant strains were classified into beneficial, deleterious, or neutral phenotypes depending on whether their relative fitness was significantly higher, significantly lower, or unchanged compared to the reference distribution.

**Table 1.** Empirical Strategy for Performance Evaluation. The testing space consists of a condition negative and condition positive datasets. The colony size datasets are generated using an isogenic population of *S. cerevisiae* grown on four 6144-density agar plates (**see Materials and Methods**). These plates were imaged at eleven time points from pinning to saturation. A subset of colonies on the plates were mocked as references, and the rest were mocked as mutants. This dataset was considered condition negative, as the reference and mutant colonies: i) are isogenic, and ii) grown to the same time point. The condition positive dataset was made up of virtual plates created by combining reference and mutant colony size data from different time points, so that the DFE is either bimodal or random. These datasets are used to measure the ability of the LI Detector to observe a variety of fitness effects. t_R_ represents the reference colony size time, and tM the mutant colony size time.

| Test Dataset | Time of Colony Size Data | Colony Size Distribution | Expected Phenotype | Performance Measure |
| --- | --- | --- | --- | --- |
| Condition Negative | $t_R = t_M$ | Uniform | Neutral | Specificity |
| Condition Positive (Virtual Plates) | $t_R > t_M$ | Bimodal | Deleterious | Sensitivity |
| | $t_R < t_M$ | | Beneficial | |
| | $t_M \geq t_R > t_M$ | Random | Deleterious<br>Neutral<br>Beneficial | |
| $t_R$ = Reference colony size time, $t_M$ = Mutant colony size time | | | | |

### Construction of condition negative and positive datasets for performance evaluation

To evaluate the performance of the LI Detector, we constructed datasets where the underlying DFE was known, but colony sizes were realistically affected by spatial biases and other technical artifacts of CBHTS (**Fig. 2A**). To this end, we applied the pin-copy-upscale experimental pipeline of our framework (**Fig. 1A**), starting with four 384-well glycerol stock plates, each containing replicate frozen cultures of the same strain (FY4^29^). This procedure generated four 6144-density agar plates containing 16 replicate colonies for each culture in the starting glycerol stock plates (**see Materials and Methods**). The sizes of these colonies were measured at eleven time points while they grew to saturation (**Fig. 2B**). The colonies originating from one of the glycerol stock plates were treated as reference, and the rest were treated as mutants.

**Figure 2.**
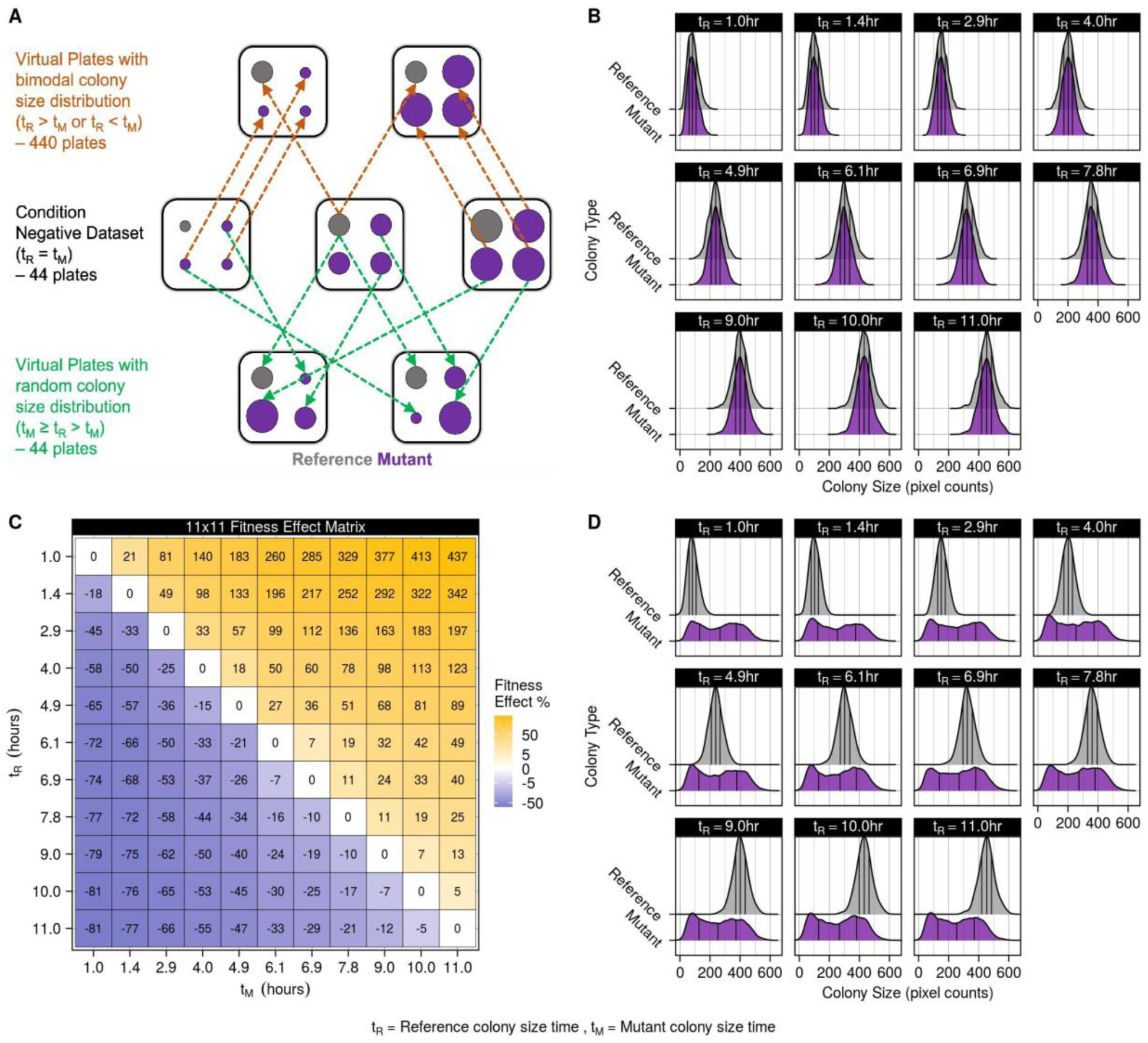
Condition negative and positive datasets used for performance evaluation. **A**. Illustration of the condition positive and negative datasets described in **Table 1**. The squares represent plates, and the circles within them represent colonies. The reference colonies are colored as grey and the mutant colonies as purple. The middle row represents the condition negative dataset shown as a single plate at three different time points. There were 44 plates in this dataset. The top row shows two virtual plates made by combining reference colony size data from one time point with mutant colony size data from another. These virtual plates had a bimodal colony size distribution. There were 440 such virtual plates. The bottom row shows two virtual plates where the reference colony size data taken from one time point is combined with mutant colony size data randomly selected from any time point. These virtual plates had a random colony size distribution. There were 44 such virtual plates. All virtual plates maintain the same spatial layout of colonies as the condition negative dataset, as is shown by the arrows. t_R_ is reference colony size time, and t_M_ is mutant colony size time. **B**. Reference and mutant population colony size density plots from the condition negative dataset. Vertical black lines within the density plots represent the lower, middle, and upper quartile. All mutants are expected to have a neutral phenotype. **C**. Fitness effect matrix of the condition positive virtual plates with bimodal colony size distribution. Mutant (t_M_) and reference colony size time (t_R_) is represented on the x-axis and y-axis, respectively. The fitness effect was calculated as the difference in mutant and reference mean colony sizes as a percentage of the reference mean colony size (**Supplementary Fig. S4**). This dataset was used to calculate the sensitivity of the LI Detector as a function of the fitness effect. **D**. Reference and mutant population colony size density plots from the condition positive virtual plates with random colony size distribution. Vertical black lines within the density plots represent the lower, middle, and upper quartile. Mutant strains could be beneficial, deleterious, or neutral. These virtual plates were used to evaluate LI Detector’s sensitivity in situations where *a priori* assumptions of fitness are challenging to make.

To estimate specificity, we assembled a “condition negative” dataset consisting of colony size measurements of our plates at eleven time points. None of the mutants in this dataset should be significantly larger (beneficial) or smaller (deleterious) than the references (**Fig. 2B**). We then assembled two artificial “condition positive” datasets consisting of “virtual plates” that we used to evaluate the sensitivity of the LI Detector (**Fig. 2C-D**). These virtual plates were constructed so that the underlying DFE would be known and readily comparable to the LI Detector and MCAT results. The first condition positive set combined colony size estimates of the mock references and mutants from two different time points, resulting in virtual plates with bimodal colony size distributions: a reference distribution, and a mutant distribution with a smaller or larger mean (**Supplementary Fig. S4**). The fitness effect was measured as the difference in the mean colony sizes of the two distributions as a percentage of the reference distribution mean colony size (**Fig. 2C**). Doing this allowed us to evaluate sensitivity for a broad range of fitness effects. The second condition positive set combined the reference distribution from a single time point with mutant colony sizes from randomly chosen time points, resulting in virtual plates with random DFE (**Fig. 2D**). The random DFE allowed us to estimate sensitivity when the traditional assumptions used for spatial bias correction are unwarranted. It is important to note that all virtual plates retain realistic spatial biases in colony sizes because they maintain the original plate layout.

We leveraged the condition negative and positive datasets to compare the performance of LI Detector (LID), LI Detector without source normalization (LID-SN), and LI Detector without local artifact correction (LID-AC) with that of MCAT^14^. We also used a random generator (RND) to assign background colony sizes by only taking the global colony size distribution of the reference population into account. Lastly, the observed colony sizes were used as-is, as “fitness” estimates to generate phenotype results when no normalization (NO NORM) was done on the datasets.

### LI Detector can accurately estimate background colony sizes and eliminate spatial bias

In our condition negative data set, variation in colony sizes should only stem from the natural biological variation expected for an isogenic population and from the spatial bias. Hence, the fitness estimates obtained after spatial bias removal should only reflect biological variability and the added noise from the bias removal process. We measured the coefficient of variance percentage (CV%) of the colony sizes and fitness estimates for images taken at multiple time points (**see Materials and Methods**). LID, LID-AC, and MCAT^14^ showed a significant reduction in CV% compared to NO-NORM, while LID-SN did not (**Supplementary Fig. S5A**). This finding indicates that the LI Detector can reduce spatial bias and confirms that source-normalization plays a vital role in doing so^14^.

The LI Detector’s ability to remove spatial bias depends on the accuracy with which it can estimate background colony sizes using the reference population colony sizes. We used the root mean square error (RMSE) between background and observed colony sizes as a percentage of the mean observed colony size to measure this. LID, LID-AC, and MCAT^14^ RMSE% were indistinguishable for the higher time points when the colonies begin to saturate (**Supplementary Fig. S5b**). RMSE% for LID-SN was significantly higher than LID (p = 0.00019, Wilcoxon rank-sum test), again indicating the importance of performing source-normalization (**Supplementary Fig. S5c**). All methods performed better than RND. Overall, these findings show the LI Detector performs as well as MCAT^14^ in eliminating spatial biases by integrating both global and local spatial contexts.

### LI Detector identifies small fitness effects with high specificity and sensitivity

To evaluate the LI Detector’s ability to detect neutral, beneficial, and deleterious fitness effects, we estimated its specificity and sensitivity using our condition negative and positive datasets, respectively (**Table 1, Fig. 2**). Specificity was calculated as the proportion of mutant strains that were correctly classified as neutral using our condition negative dataset (**Fig. 2B**). LID’s specificity was above 98% for an empirical p-value cut off of 0.05 and remained above 95% when that cut-off was increased to an empirical p-value of 0.1. For comparison, MCAT^14^ showed a maximum specificity of 94.5% for an empirical p-value cut off of 0.05 using the same dataset (**Fig. 3A**).

**Figure 3.**
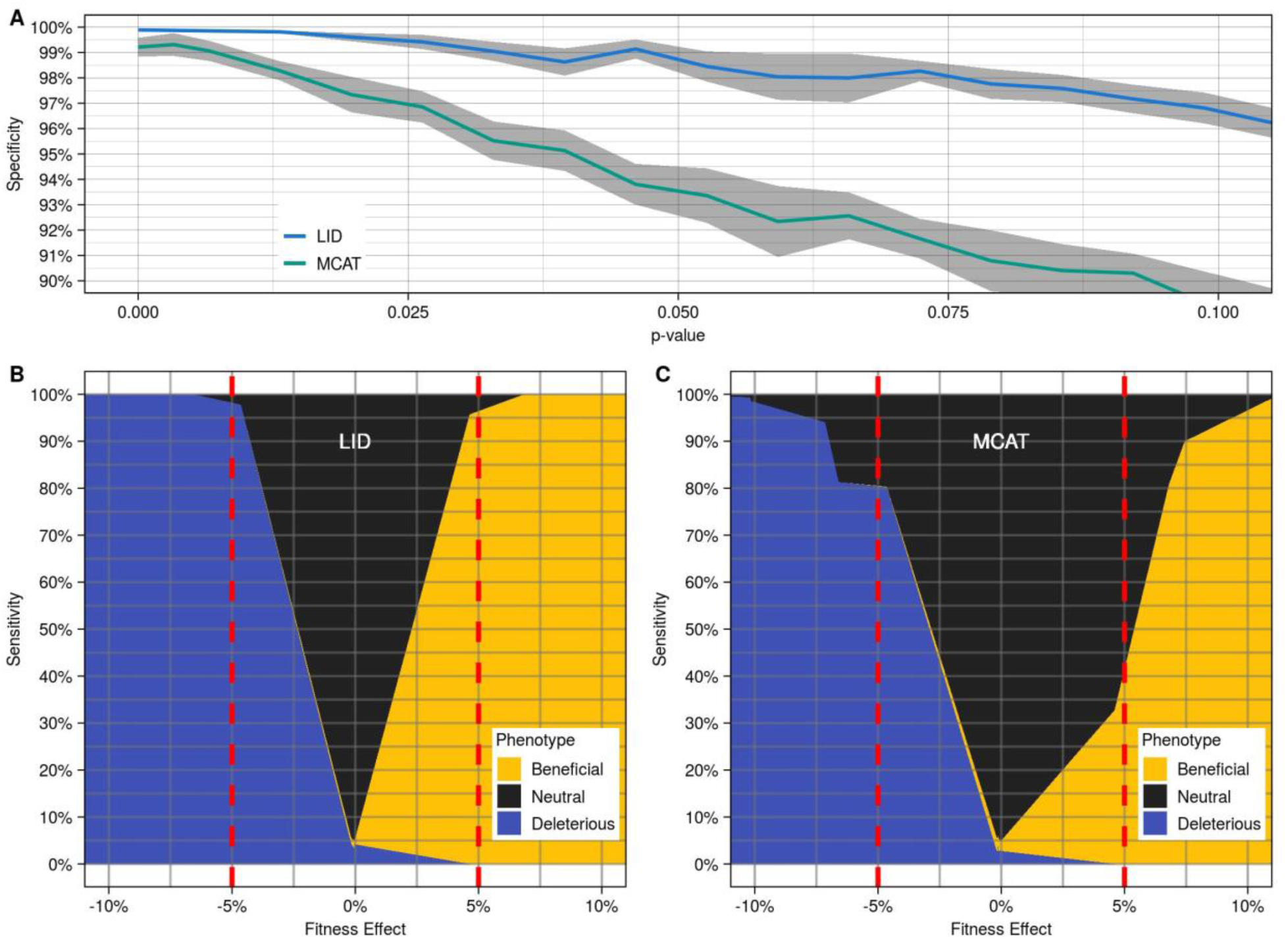
The LI Detector has high specificity and sensitivity. **A**. Average specificity (solid colored line) and standard error (gray region) at various empirical p-value cut-offs for LID (blue) and MCAT^14^ (green). Empirical p-values (x-axis) calculated using the reference strain relative fitness distribution (**see Materials and Methods**). Specificity (y-axis) was estimated using the condition negative dataset as the proportion of mutants classified as neutral (**see Materials and Methods**). **B**. LID phenotype classification results from the virtual plates with bimodal distribution are arranged according to increasing fitness effects. Here, the fitness effect is the mean mutant and mean reference colony size difference as a percentage of the reference colony size for each virtual plate. Sensitivity is calculated as the proportion of mutants correctly identified as significantly different (beneficial or deleterious) than the reference for each fitness effect value. The dotted red line indicates a 5% fitness effect. A 5% false positive rate was maintained while generating these results. **C**. MCAT^14^ phenotype classification results from the same data as **B**.

Sensitivity was estimated as the proportion of mock mutant strains correctly classified as either beneficial or deleterious at a false positive rate of 5% using our condition positive dataset with bimodal fitness distribution (**Fig. 2C, Supplementary Fig. S4**). LID’s sensitivity was higher than 95% for beneficial and deleterious fitness effects of 5%, reaching 100% for fitness effects of about 7% (**Fig. 3B**). These findings show that LID is highly sensitive in observing small fitness effects; notably, it is equally sensitive to increases and decreases in fitness. This result depended on the fitness estimation strategy, with LID performing significantly better than LID-AC, LID-SN, and NO-NORM (**Supplementary Fig. S6**). We also performed the same analysis using MCAT^14^. MCAT^14^ was 80% sensitive in detecting 5% fitness decreases, and only 40% sensitive when it came to 5% fitness increases (**Fig. 3C**). We hypothesize that MCAT’s lower sensitivity stems from its use of a local window of surrounding mutants rather than a reference colony grid to estimate background colony size. These results show that the LI Detector displays improved sensitivity, remarkably so for beneficial effects, for the same specificity as MCAT.

### LI Detector maintains high sensitivity when the DFE is random

We designed the LI Detector to be highly sensitive regardless of the underlying DFE. To evaluate LI Detector’s performance when the underlying DFE is random, we used our condition positive data set made of forty-four virtual plates with random colony size distribution (**Fig. 2D)**. The forty-four plates combined contained 41.60% beneficial and 50.26% deleterious mutants with sixteen replicate colonies of each (**Fig. 4A**). We found that LID was 98.93% sensitive, successfully identifying 98.65% of beneficial and 99.20% of deleterious mutants (**Fig. 4B**). In comparison, MCAT^14^ was 83.08% sensitive and successful in identifying 82.76% of beneficial and 83.40% of deleterious mutants (**Fig. 4C**). The false positive rate was maintained at 5% for both methods. Virtual plate-wise phenotype classification results show that the actual classification (**Fig. 4D**) is better captured by LID (**Fig. 4E**) and that MCAT^14^, in general, had more false negatives (**Fig. 4F**). LID’s neutral calls were mostly limited to fitness effects of 5% or smaller, whereas MCAT^14^ neutral calls covered a wider range of fitness effects (**Fig. 4G-I**). That MCAT^14^ was considerably less sensitive than LID in this scenario was not surprising, since a random underlying distribution of fitness effects violates the assumptions of MCAT^14^ and other existing methods.

**Figure 4.**
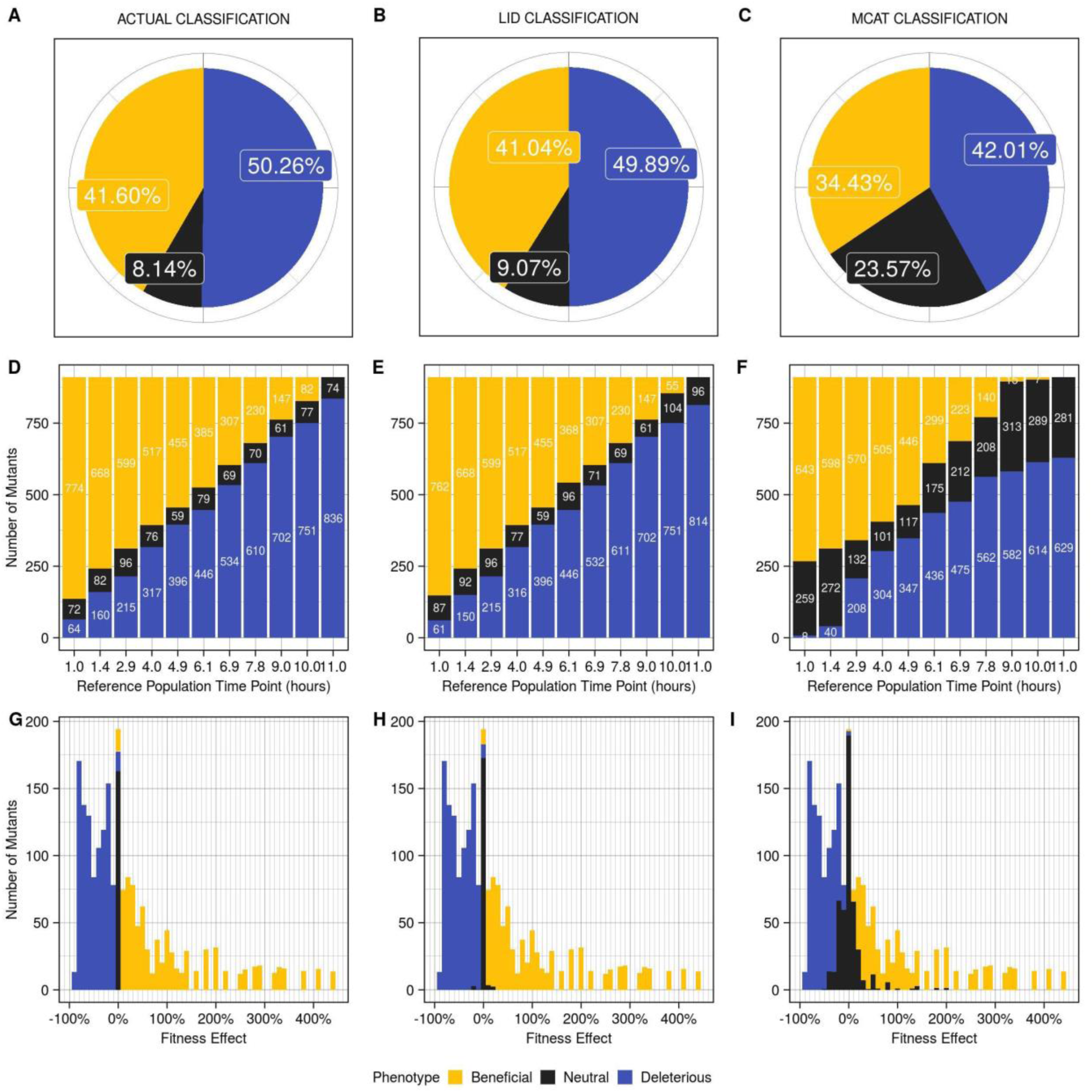
The LI Detector maintains high sensitivity even when the underlying DFE is random. **A**. The actual classification of the mutants in the random DFE condition positive dataset, per construction, with 41.60% beneficial, 50.26% deleterious, and 8.14% neutral. **B**. and **C**. show the classification results from LID and MCAT^14^, respectively. **D-F**. Bar graphs showing **D**. actual, **E**. LID, and **F**. MCAT^14^ classification of mutants for each virtual plate with random DFE. The virtual plates are arranged according to their reference colony time point. **G-I**. Bar graph of pooled results from all plates arranged according to the fitness effects for the **G**. actual classification, **H**. LID and **I**. MCAT^14^. Each bar has a width of 10%. False positive rate was maintained at 5% for both LID and MCAT^14^ in these analyses.

### The LI Detector’s sensitivity increases with an increasing number of references and replicates

LI Detector’s superior performance comes at the cost of having to integrate a reference colony grid, and therefore use a higher number of plates to screen the same number of mutant colonies. We analyzed how the number of references per plate and the number of replicates per strain affected LID’s sensitivity. To do this, we computationally masked portions of the reference colony grid and replicates, and then reanalyzed the virtual plates with bimodal and random DFE in our condition positive dataset (**see Materials and Methods**). We observed that LID’s sensitivity in detecting 5% fitness effects increased in proportion to the number of reference colonies per plate, as well as to the number of replicates per strain in both sets of virtual plates (**Fig. 5**). Unsurprisingly, sensitivity was higher for detecting a fitness effect of 7% (**Supplementary Fig. S7A**). Increasing the number of replicates was most powerful when there were more references on the plate (**Supplementary Fig. S7B**). In general, the sensitivity was higher in the virtual plates with bimodal than random DFE (**Fig. 5**). These observations are consistent with the finding that RMSE% is inversely related to the number of reference colonies per plate (**Supplementary Fig. S8**). On the other hand, LID’s specificity was consistently above 95%, independent of the fitness estimation strategy (**see Materials and Methods**), the proportion of references per plate, and the number of replicates per mutant strain (**Supplementary Fig. S9**). The LI Detector users may choose the number of references and replicates adequate for their purposes as a function of the fitness effects they expect to observe and the sensitivity they aim to achieve.

**Figure 5.**
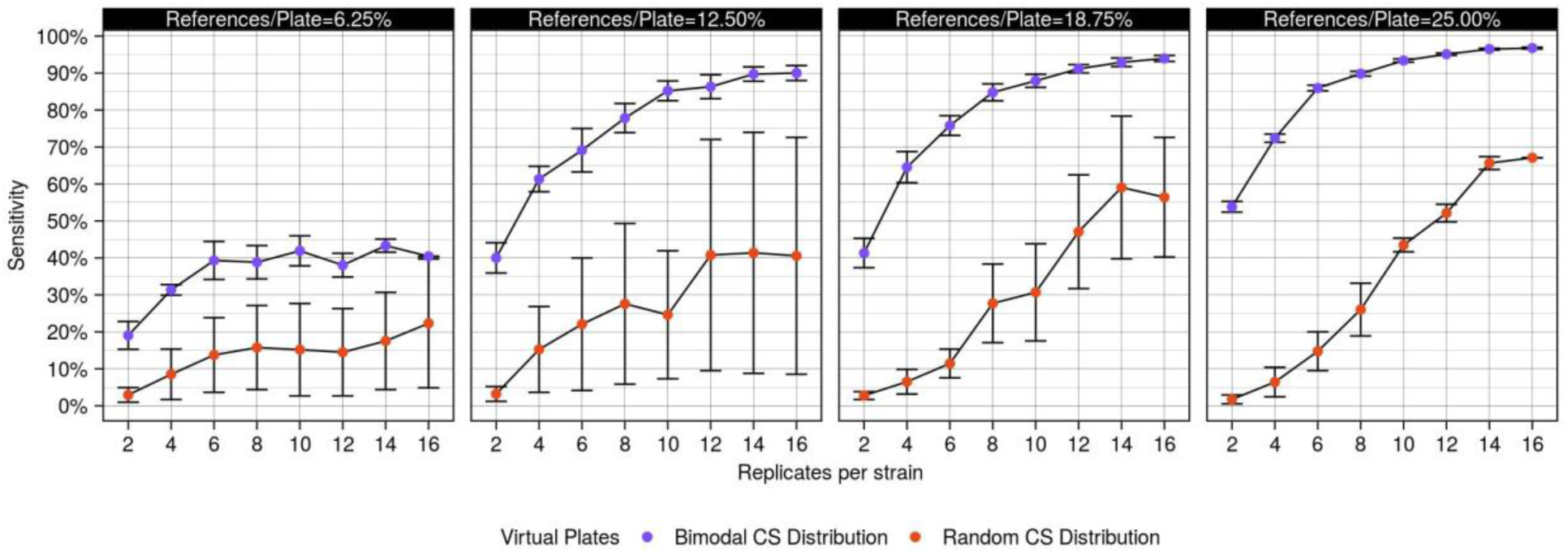
Sensitivity is directly related to the number of references and replicates. Sensitivity for observing 5% fitness effects, as a function of the varying proportion of references per plate (individual panels) and the number of replicates per strain (x-axis) was estimated for virtual plates with bimodal (purple) and random (orange) colony size distribution. Error bars represent a single standard deviation.

## DISCUSSION

LI Detector is a CBHTS framework (**Fig. 1**) that generates reliable and well-resolved fitness estimations without being dependent on *a priori* assumptions of the DFE (**Fig. 3A-B, Fig. 4B, E, H**). LI Detector is specifically designed to observe small deleterious and beneficial changes in fitness (**Fig. 3B**). Therefore, it is a valuable method for improving the current gene-gene, gene-environment and protein-protein interaction networks for colony-forming-microorganisms.

Existing spatial bias correction methods work best in unbiased genome-wide studies with a large number of plates and mutants^17^. While alternate methods have been developed to increase sensitivity at the small scale level^24^, LI Detector provides a flexible approach that can be applied to CBHTS independent of their scale and of the choice of strains to screen. For example, LI Detector can be used as efficiently for a highly biased screen of non-synonymous mutations in a single gene to identify important residues^30-36^, or for a genome-wide synthetic genetic array used to infer genetic interactions^6,37-40^. This freedom of experimental design expands the applicability of CBHTS for broader scientific inquiry.

We show that LI Detector has the power to uncover significant fitness effects as small as 5% with 95% sensitivity when 25% of the plate is dedicated to reference colonies and mutant strains are represented by 16 replicates colonies (**Fig. 3b, Fig. 5**). Smaller fitness effects can be observed with comparable sensitivity by increasing the number of replicates per strain (**Supplementary Fig. S7A**). Existing methods, like MCAT^14^, also provide quantitative output of fitness with high resolution; however, without a reference grid and proper spatial bias correction, one cannot statistically determine if the small effects are meaningful. LI Detector’s ability to detect small increases in fitness, in particular, makes it a favorable method to examine gain-of-function mutations, questions of evolutionary biology, and pharmacological screens of adaptation and resistance^10,28,41-48^. The unprecedented sensitivity of LI Detector augments the discovery potential of CBHTS.

A caveat of LI Detector is that a portion of the colony positions on the plates is sacrificed for reference colonies that could otherwise be used for mutants. Consequently, this increases the overall resources required for the experiment, including media, number of plates, storage space, pinning time, and imaging time. We have shown that LI Detector’s accuracy in predicting background colony sizes and its sensitivity in detecting small fitness effects is directly related to the proportion of reference colonies on a plate (**Fig. 5**). However, the proportion of references per plate and the number of replicates per strain can be tunable according to the user’s requirement. It must be noted that the cost of reducing the number of references is lower for detecting more substantial fitness effects. For example, sacrificing 12.5% of the plate for reference colonies instead of 25% has almost no detriment to detecting 7% fitness effects (**Fig. 5, Supplementary Fig. S7a**). A higher number of references and replicates can be used if the goal is to look for minute changes in fitness, as are frequently observed with the deletion of non-essential genes or minor changes to the coding sequence of a given gene. Alternatively, fewer references and replicates may be used where larger fitness effects are expected or desired, such as finding the most drug-resistant mutant. That said, users interested in large fitness effects exclusively may use existing methods like MCAT^14^ instead of the LI Detector to save resources, as long as *a priori* assumptions of the DFE are reasonable to make.

In summary, the LI Detector framework experimentally introduces a reference population grid on plates whose colony size estimates are used to correct for spatial bias independently of the underlying DFE. It has the potential to expand the utility of CBHTS by making them independent of scale, sensitive towards small fitness effects, and equally sensitive in detecting increases and decreases in fitness. Although developed and validated using *S. cerevisiae*, it can be applied to any colony-forming-microorganisms, including clinically relevant isolates, as long as they can be grown in the laboratory.

## MATERIALS AND METHODS

### Validation experiment using an isogenic population

A method validation experiment was conducted using an isogenic population that was mocked as either references or mutants.

#### Yeast strain, medium and robotic equipment

A previously characterized prototrophic *S. cerevisiae* strain in the S288C background, FY4^29^, was used to conduct experiments in YPD medium (1% w/v yeast extract, 2% w/v peptone, 2% w/v dextrose, and 2% agar in the case of solid medium). A single colony of FY4^29^ selected from a streak out was used to inoculate liquid YPD medium and grown overnight at 30 **°**C. This culture was used to create four 384-well glycerol stocks with wells containing 18μL of 50% glycerol and 42μL of culture media. Two to five wells in each stock were left empty to create gaps in the colony grid when pinned on solid medium. The stocks were stored at -80**°**C before use. The benchtop RoToR HDA robotic plate handler (Singer Instruments Co Ltd, Roadwater, UK) was used for plate to plate cell transfer.

#### Pin-copy-upscale

The LI Detector experimental pipeline follows a pin-copy-upscale protocol when starting from frozen glycerol stocks. The copy-upscale steps are repeated until the desired colony density is reached (**Fig. 1A**). The four glycerol stock plates were pinned at 384-density to generate working copy agar plates. This process was performed using the RoToR HDA robot with default settings (**Supplementary Table S2**). The working copies were incubated at 30**°**C for 60 hours to reach saturation. These were then copied 1-to-1 to make transition plates (#1) using default RoToR HDA settings (**Supplementary Table S2**) and incubated at 30**°**C for 48 hours. Distinct combinations of the four transition plates (#1) were then condensed to make four 1536-density upscale plates (#1) using default RoToR HDA settings (**Supplementary Table S2**). The distinct combinations ensure that colony grids from each plate occupy different positions on the four higher-density plates. The upscale plates (#1) were incubated at 30**°**C for 30 hours, after which they were copied 1-to-1 to transition plates (#2) using custom RoToR HDA settings (**Supplementary Table S2**). The overshoot setting value at the target plate was increased to compensate for the agar surface’s unevenness and the smaller pin size of the higher density pin pads. These plates were incubated at 30**°**C for 30 hours. Four upscale plates (#2) were then made by condensing the four transition plates (#2) in distinct combinations using default RoToR HDA settings (**Supplementary Table S2**). These were incubated at 30**°**C until they reached saturation and imaged at the following eleven time points: 1.0, 1.4, 2.9, 4.0, 4.9, 6.1, 6.9, 7.8, 9.0, 10.0, 11.0 hours. All images are available at https://pitt.box.com/s/xbchjoa4ta3oq2g50q4avfypjrgz7poq.

For the purposes of evaluating the performance of LI Detector, colonies originating from a random working copy were mocked as reference strains, while the colonies from the other three working copies were mocked as mutant strains. In the upscale plates (#2) used for our analyses, 1/4^th^ of all colonies correspond to references, and the rest are treated as mutants. These plates had 16 technical replicates for every colony that was present in the working copy. **Supplementary Figure S10** provides a simplified visual representation of the plates at all pinning stages.

### Colony size estimation

Raw estimates of colony sizes are an input to the LI Detector framework (**Fig. 1B**) and can be obtained in the user’s manner of choice. Here, a custom-made lightbox with an overhead camera mount was built to acquire high-resolution images using a commercially available SLR camera (18Mpixel Rebel T6, Canon USA Inc., Melville, NY, USA). The 6144-density upscale plates (#2) were imaged at eleven time points beginning right after pinning until the colonies reached saturation, around 11 hours later. Saturation was determined as the point at which the colonies would touch each other if the plates were incubated for any longer. The images were analyzed in bulk using the “analyze_directory_of_images()” function of the MATLAB Colony Analyzer Toolkit (MCAT)^14^ with the default threshold parameter (1.25) to provide colony size estimations (https://github.com/sauriiiin/lidetector/blob/master/imageanalyzer.m). The output files containing colony size information along with the images is available at https://pitt.box.com/s/xbchjoa4ta3oq2g50q4avfypjrgz7poq.

### Spatially cognizant colony size database

A unique position identifier (pos) was given to every possible colony position across the different plates of the experiment. Each pos was linked to plate density, plate number, column number, row number and stored in a “position to coordinate” table (pos2coor). A “position to mutant name” table (pos2orf_name) was used to store information on which colony position was occupied by which mutant. The colony size estimations, along with the pos2coor table, were used to store the colony sizes in a spatially cognizant manner. **Supplementary Figure S10** is a visual representation of the plate maps made using the pos2coor and pos2orf_name tables. The colonies’ spatial layout and identity are an input to the LI Detector framework (**Fig. 1B**) and should be provided in this format by users. The format, along with the data collected for this manuscript, is available at https://github.com/sauriiiin/lidetector.

### LI Detector analytical pipeline

The LI Detector analytical pipeline (**Fig. 1B**, https://github.com/sauriiiin/lidetector/blob/master/lid.m) is designed to make fitness assessments using local reference colony information.

#### Border colony removal

Border colonies tend to grow larger because of increased access to nutrients^14-17,23^. To remove this artifact, we ignore colony size estimations of one, two, and four border rows and columns from 384, 1536, and 6144-density plates, respectively. Doing this resulted in 4864 colonies for 304 mock references and 14576 colonies for 911 mock mutants across four 6144-density plates. All further analysis is done using this set.

#### Local artifact correction (AC)

Local artifact correction is inspired by the “competition correction” feature present in existing tools^15,16^. An “artifact score” is assigned to every colony on a plate as a ratio of its colony size compared to its current and past neighbors. The current neighbors are a colony’s eight immediate neighbor colonies, and the past neighbors are eight neighboring colonies that were pinned from the same source plate. The reference population’s artifact scores are used to determine outliers, defined as two median adjusted deviations or more from the median. Outliers are defined as colonies growing disproportionately big or small as compared to their neighboring colonies. Outliers that occur as a localized group of three or more neighbors of both big and small colonies are considered for correction. The less abundant outlier in the group is expected to have driven the phenotype. For example, a single small or dead colony would increase the relative access to nutrients for all its neighbors, which would all be expected to grow bigger than u sual and vice-versa. Raw colony sizes of all the driver’s immediate neighbors are median normalized using the median reference population colony size for the plate. Users have the option to skip this correction.

#### Source normalization (SN)

LI Detector uses a source-based computational deconstruction of high-density plates into their four lower-density sources to correct the source-related colony size differences introduced during the upscaling process (**Supplementary Fig. S2**). This correction is a reimplementation of MCAT’s interleave filter^14^. Each source-deconstruct is individually normalized in the later steps, making it necessary for the penultimate density plates to have a reference population grid. Users have the option to skip this correction, although we strongly recommend against skipping if upscales are performed.

#### Reference-based normalization

A two-dimensional linear interpolant is applied to the reference population grid to estimate expected colony sizes on the entire colony grid. This reference colony based estimated colony size is referred to as the “background colony size.” The background colony sizes represent the predicted reference colony growth on every position of the grid conditioned upon the spatial context. Relative fitness is estimated as the ratio of the local artifact corrected colony size to the background colony size, thus controlling for spatial context.

### Different strategies for fitness estimation

The LI Detector analytical pipeline is applied to colony size estimates to control for spatial bias and measure relative fitness as described above. The analytical pipeline is used as-is (LID), without local artifact correction (LID-AC), and without source-normalization (LID-SN) to measure the impact of these components on the downstream analysis. Raw observed colony size estimates were also used as “fitness” measurements without performing any normalization (NO-NORM). Fitness estimates were also made using the MCAT’s^14^ SpatialMedian normalization with window size nine along with the Interleave filter (https://github.com/sauriiiin/sau-matlab-toolkit/blob/master/image2resBEAN.m).

### Measuring spatial bias and the accuracy of background colony size

The coefficient of variance of fitness and colony size estimations was used to measure the impact of spatial bias in colony sizes of an isogenic population (**Supplementary Fig. S5A**). Ten random observations were picked, with replacement, 2000 times to measure the coefficient of variance as a percentage of the mean (CV%). CV% distributions for LID, LID-AC, LID-SN, MCAT^14^, and NO-NORM were compared using the Wilcoxon rank-sum test.

The accuracy of background colony size was measured using root mean square error (RMSE) estimation as a percentage of the average observed colony size (**Supplementary Fig. S5B**). A random colony size predictor (RND) was used as a null model for background colony size prediction. The RND generated random colony sizes from a normal distribution, with the *rnorm* function in R^49^ using the mean and standard deviation of observed colony sizes. The Wilcoxon rank-sum test was used to compare RMSE results from LID, LID-AC, LID-SN, MCAT^14^, and RND.

### Calculating significant fitness changes and assigning phenotypes

The relative fitness of each strain was measured as the mean of estimated relative fitness among its replicates. This measurement was done after removing the outlier observations based on three median adjusted deviations. The reference strain relative fitness distribution was used as a null distribution for hypothesis testing, as the reference strains are isogenic, and no real fitness differences are expected. An empirical p-value was estimated for all mutant strains based on where they fall relative to this null distribution (https://github.com/sauriiiin/lidetector/blob/master/lid.m). For example, an empirical p-value of

0.05 or below would mean that the mutant’s relative fitness is in the top or the bottom 2.5^th^ percentile of the reference fitness distribution. The phenotype of mutant strains significantly different from the reference population is classified as “beneficial” or “deleterious,” depending on whether its estimated relative fitness is above or below 1. The remaining mutant strains that do not have a significant change in fitness are classified as having a “neutral” phenotype.

### Empirical strategy for performance evaluation

An empirical strategy was devised to thoroughly examine the LI Detector’s performance. A condition negative and positive dataset were created to estimate specificity and sensitivity, respectively (**Table 1, Fig. 2A**).

The condition negative dataset consisted of data where the mock mutants and references have similar colony size distribution. To this end, colony size data taken from any time point represents a unique condition negative dataset (**Fig. 2B**). We tested 44 such plates, four plates for the 11 time points that images were taken. The proportion of mock mutant strains that are successfully called neutral by the LI Detector represents the true negative rate or specificity.

The condition positive dataset consisted of colony size data where mutant strains can be deleterious or beneficial. Two sets of virtual plates were created to generate such a condition positive dataset. The first set of virtual plates contained a bimodal distribution of colony sizes (**Supplementary Fig. S4, S11**) where colony size estimations for reference and mutant colony positions came from two different time points while maintaining their topological context (https://github.com/sauriiiin/paris/blob/master/techPowA.m). The fitness effect between the reference and mutant colony size distribution is the difference of their mean colony sizes as a percentage of the mean reference colony size (**Fig. 2C**). We tested 440 virtual plates with bimodal colony size distribution resulting from combining reference colony size data from 11 time points (t_R_) with mutant colony size data taken from 10 time points (t_M_) and having four plates for each t_R_ - t_M_ combination.

The second set of virtual plates contained a random distribution of colony sizes were reference colony size data from a particular time point was combined with mutant colony size data randomly selected from all time points (**Fig. 2D**). Colony size estimates for replicates of the same mutant were all selected from the same time point (https://github.com/sauriiiin/adaptivefitness/blob/master/scripts/4CX/4C_MESSUP.R). We tested 44 virtual plates with random colony size distribution by having 4 plates for the 11 time points that reference colony size data (t_R_) can be taken from (**Fig. 2D**).

Mutants that are successfully called beneficial or deleterious in these virtual plates are used to estimate the true positive rate or sensitivity of the LI Detector (https://github.com/sauriiiin/adaptivefitness/blob/master/scripts/4CX/4C_POWDY.R). For the virtual plates, an empirical p-value cut off that controls the false positive rate at 5% was used to make the examination of sensitivity comparable between LID, LID-AC, LID-SN, MCAT^14^, and NO-NORM. The results from this analysis are represented in **Fig. 3, Fig. 4, Supplementary Fig. S6**, and **Supplementary Fig. S9**.

### Measuring the impact of the number of references and replicates

The reference colony proportion was sequentially reduced from 25% to 18.75% to 12.5% to 6.25% by masking 1/4^th^ of the existing reference grid each time. This reduction was achieved by masking colonies on the 384-density mock reference plate and then propagating those masked colonies through the other densities. The number of replicates per strain was reduced in parallel by masking the n^th^ replicate of every mock strain. Replicates were reduced from 16 to 2 in increments of 2. This process was repeated ten times to mask a variety of replicate combinations (https://github.com/sauriiiin/paris/blob/master/techPowA.m). Results from the analysis of the resultant plates are represented in **Figure 5** and **Supplementary Figure S7A-B**.

## Data availability

All data generated/analyzed in this study is available in the main text, in the Supplementary Figures and Tables, and as Supplementary Data files. All supplementary data are also on GitHub: https://github.com/sauriiiin/lidetector.

## Code availability

The code is available to download at https://github.com/sauriiiin/lidetector, along with instructions on how to use it. Image processing, relative fitness estimations, and analyses presented in the result section are available at https://github.com/sauriiiin/sau-matlab-toolkit. All images within the main article and supplementary data were generated using code available at https://github.com/sauriiiin/adaptivefitness/tree/master/scripts/paper.

## ACKNOWLEDGEMENTS

The authors are grateful: Dr. Aaron Wacholder for discussions; and to Dr. Branden Van Oss, Dr. Aaron Wacholder, Kate McCourt, Tyler Fortuna, and Osama Shiraz Shah for reviewing the manuscript prior to submission. This work was supported by: funds provided by the Searle Scholars Program to A.-R.C.; the National Institute of General Medical Sciences of the National Institutes of Health grants R00GM108865 and DP2GM137422 awarded to A.-R.C.

## AUTHOR CONTRIBUTIONS

Conceptualization: S.B.P., A.-R.C.; Methodology: S.B.P., A.-R.C.; Investigation: S.B.P., N.C.C.; Writing-original draft: S.B.P.; Writing-review and editing: S.B.P., A.-R.C., N.C.C.; Supervision: A.-R.C.

## SUPPLEMENTARY INFORMATION

**Table S1.** Existing bioinformatics tools for colony-based screen image analysis and normalization.

| Tool | Experimental Design | User Input | Normalization Options | Output | Reference # |
| --- | --- | --- | --- | --- | --- |
| HT Colony Grid Analyzer | Synthetic genetic arrays (SGA), Epistatic miniarray profile (E-MAP) | 96 to 1536 density plate images | Plate middle mean (PMM) normalization | Averaged S Scores | 21 |
|  |  | Assign top left colony position | Filter strain based on noise and linkage |  |  |
| Colonyzer | Any rectangular array format | 48 to 1536 density in rectangular or circular petri dish | Spatial lighting gradient | Raw colony size | 18 |
|  |  |  | Median background intensity correction | Integrated optical density (IOD) |  |
|  |  |  |  | Colony color |  |
|  |  |  |  | Colony granularity |  |
| Balony | Any plate-based array | Single or composite images | Plate median correction | Raw colony size | 16 |
|  |  |  | Row/Column | Normalized colony size |  |
|  |  |  | Spatial correction for media thickness | Genetic interaction results |  |
|  |  |  | Competition |  |  |
|  |  |  | Filter strain based on linkage (for SGA) |  |  |
| ScreenMill | SGA | 384 to 1536 single or composite image | Artifact removal | Raw colony size | 19 |
|  |  |  | Plate median correction | Growth ratio, z-value, p-value, position data |  |
| Spotsizer | Any plate-based array | Plate images |  | Raw colony size | 50 |
| Gitter | Any plate-based array | Plate images |  | Raw colony size | 51 |
| SGAtools | Any plate-based array | 96 to 1536 density plate images | Plate middle mean (PMM) normalization | Raw colony size | 15 |
|  |  |  | Row/Column | Normalized colony size |  |
|  |  |  | Spatial correction for media thickness | Genetic interaction results |  |
|  |  |  | Competition | GO enrichment |  |
|  |  |  | Batch |  |  |
|  |  |  | Filter for large colonies and linkage (for SGA) |  |  |
| Scan-o-matic | Time lapsed plate-based array | Composite time-lapsed images from custom setup | Light/color correction | Raw colony size | 23 |
|  |  |  | Population size estimation | Population size estimates |  |
|  |  |  | Growth curve smoothing | Growth parameters |  |
|  |  |  | Reference surface-based doubling time normalization |  |  |
| MATLAB Colony Analyzer Toolkit | Any plate-based array | 96 to 24576 density plate images | Plate mode correction | Raw colony size | 14 |
|  |  |  | Edge correction | Normalized colony size |  |
|  |  |  | Source-based normalization | Colony color |  |
|  |  |  | Local median filter | Pixel intensity |  |
| LI Detector | Any plate-based array | 96 to 24576 density plate images | Local artifact | Local artifact corrected colony size | Current study |
|  |  |  | Source-based normalization | Reference-based background colony size |  |
|  |  |  | Local reference-based normalization | Relative fitness |  |

**Table S2.**
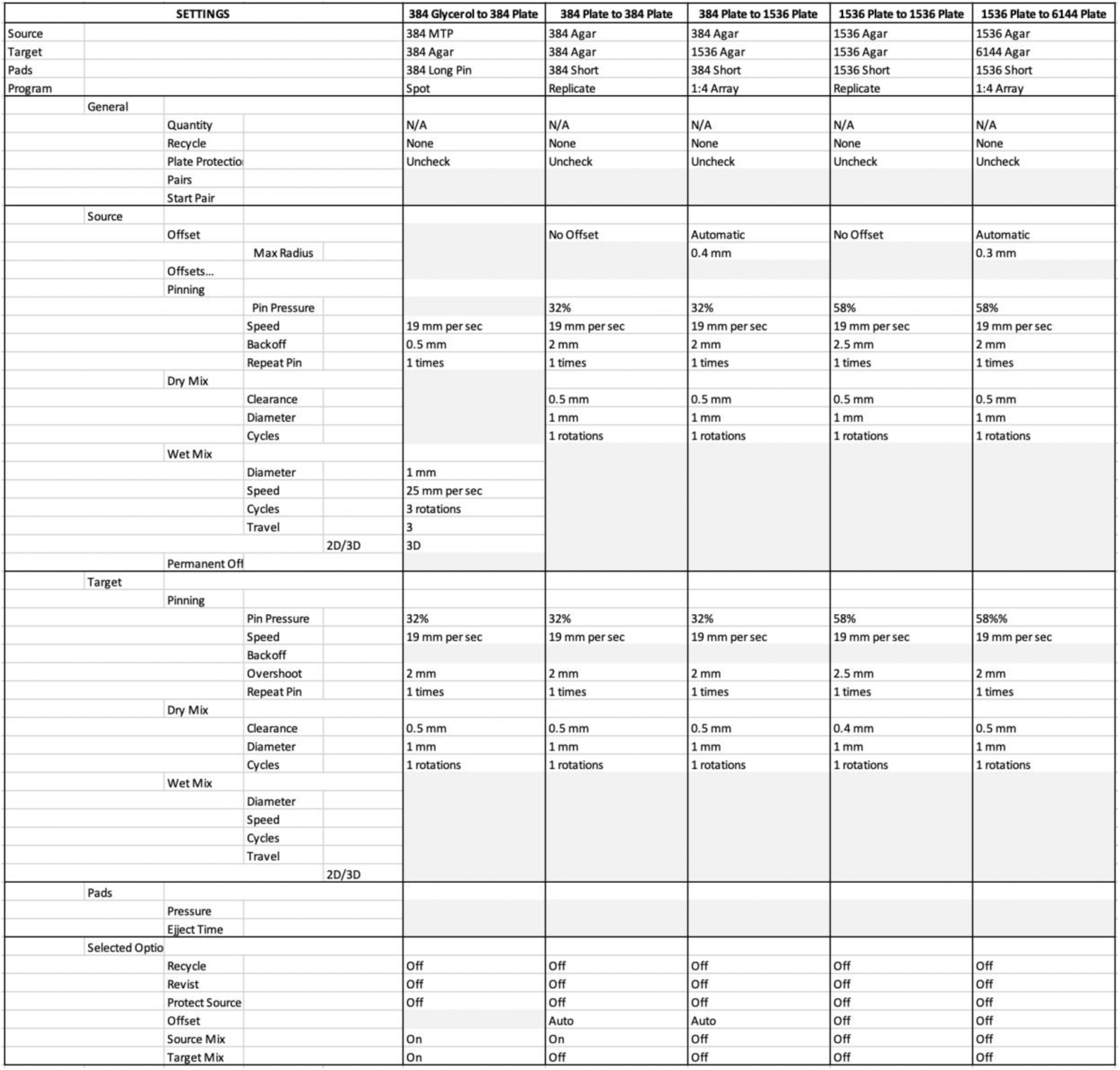
Singer RoToR settings used for the experimental pipeline.

**Figure S1.**
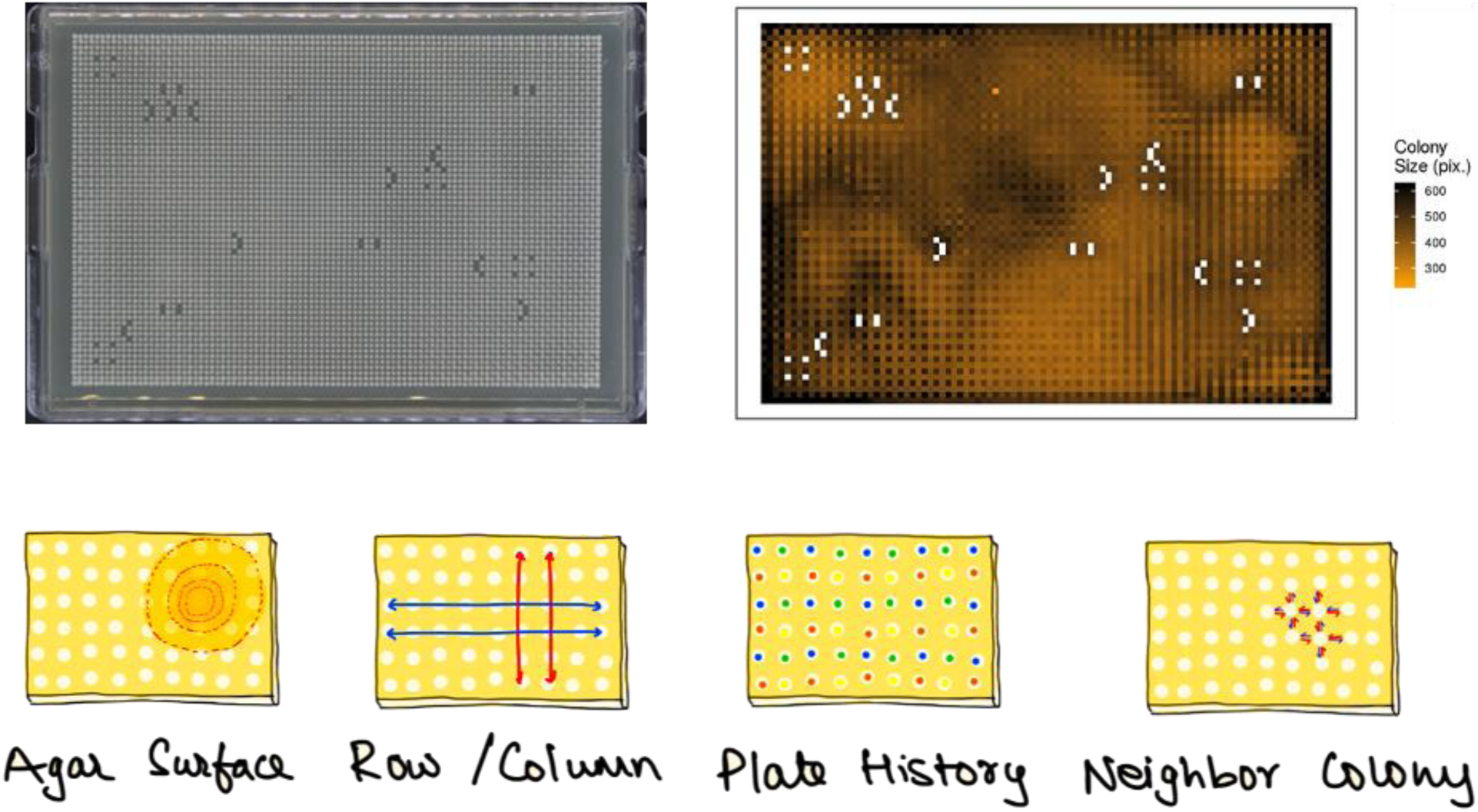
Spatial bias on a 6144-density plate. The image on the top left is a 6144-density plate having an isogenic population. The top right shows a heatmap of the same plate where each tile represents a colony and is colored according to its colony size estimation (pixel counts), going from brown (low) to black (high). On the bottom is an illustration of the types of spatial biases expected on a high-density plate.

**Figure S2.**
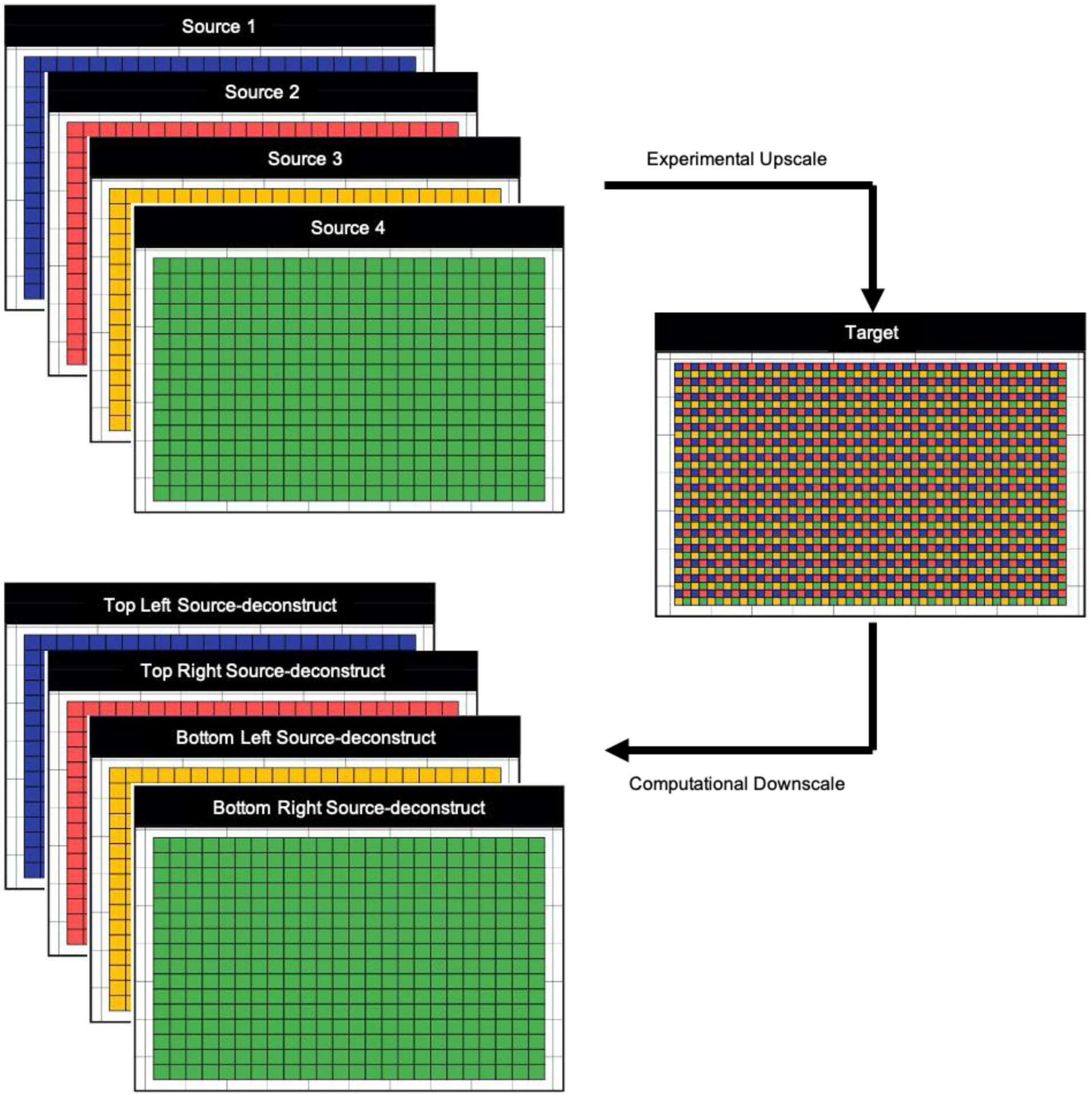
Source normalization step of the LI Detector. The target plate on the right represents a high-density plate made by experimentally condensing four lower density sources on the top left. A different color represents each source, and tiles represent colonies. This step introduces a systematic source-based bias in colony sizes that needs to be corrected. LI Detector implements a source normalization (SN) step, where it computationally downscales the colony size estimations of the higher density plate into its four-corresponding source-deconstructs shown on the bottom left. These source-deconstructs are individually normalized during the downstream analysis.

**Figure S3.**
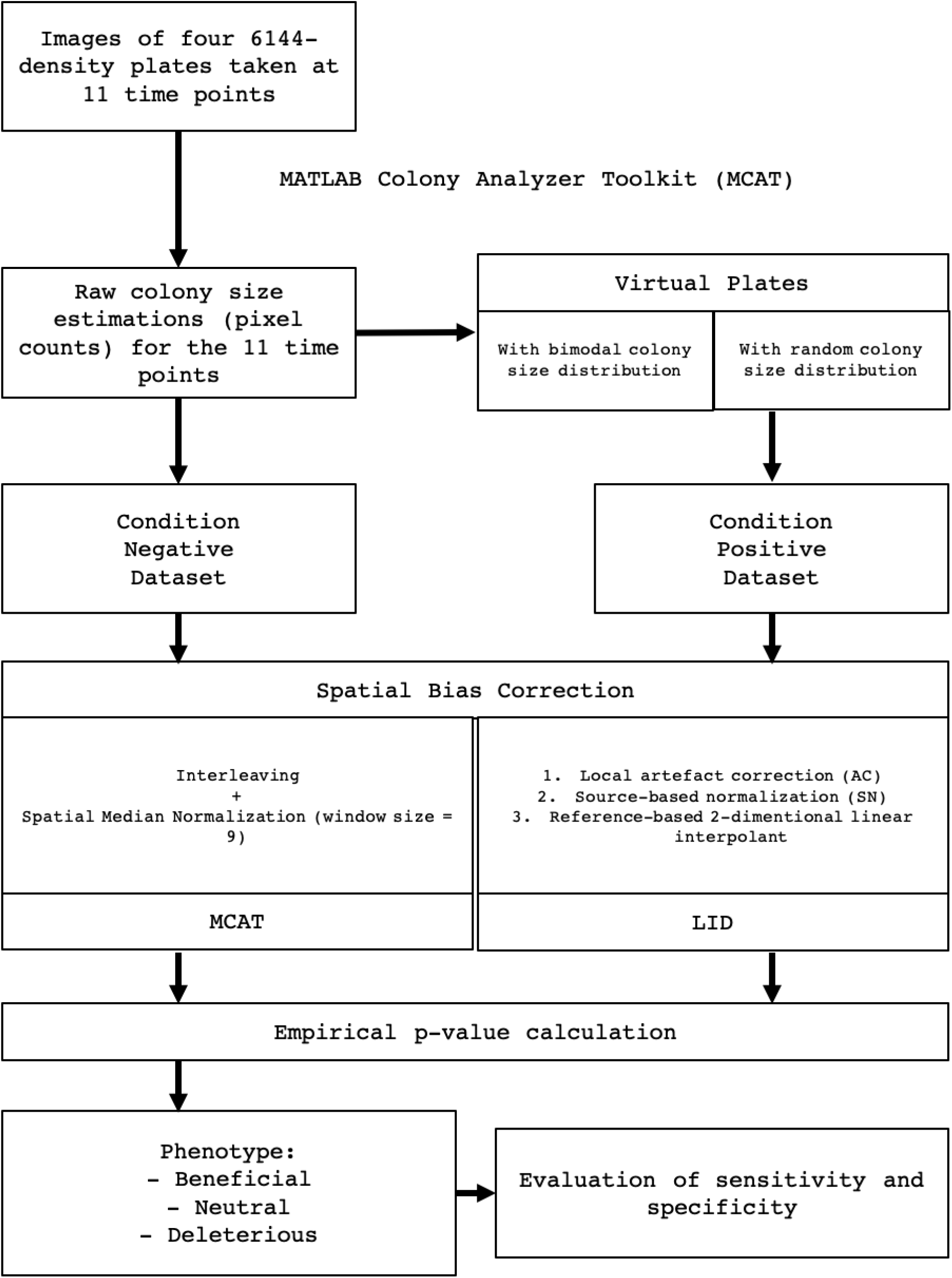
Workflow of the methodology adopted for analyzing LI Detector’s performance.

**Figure S4.**
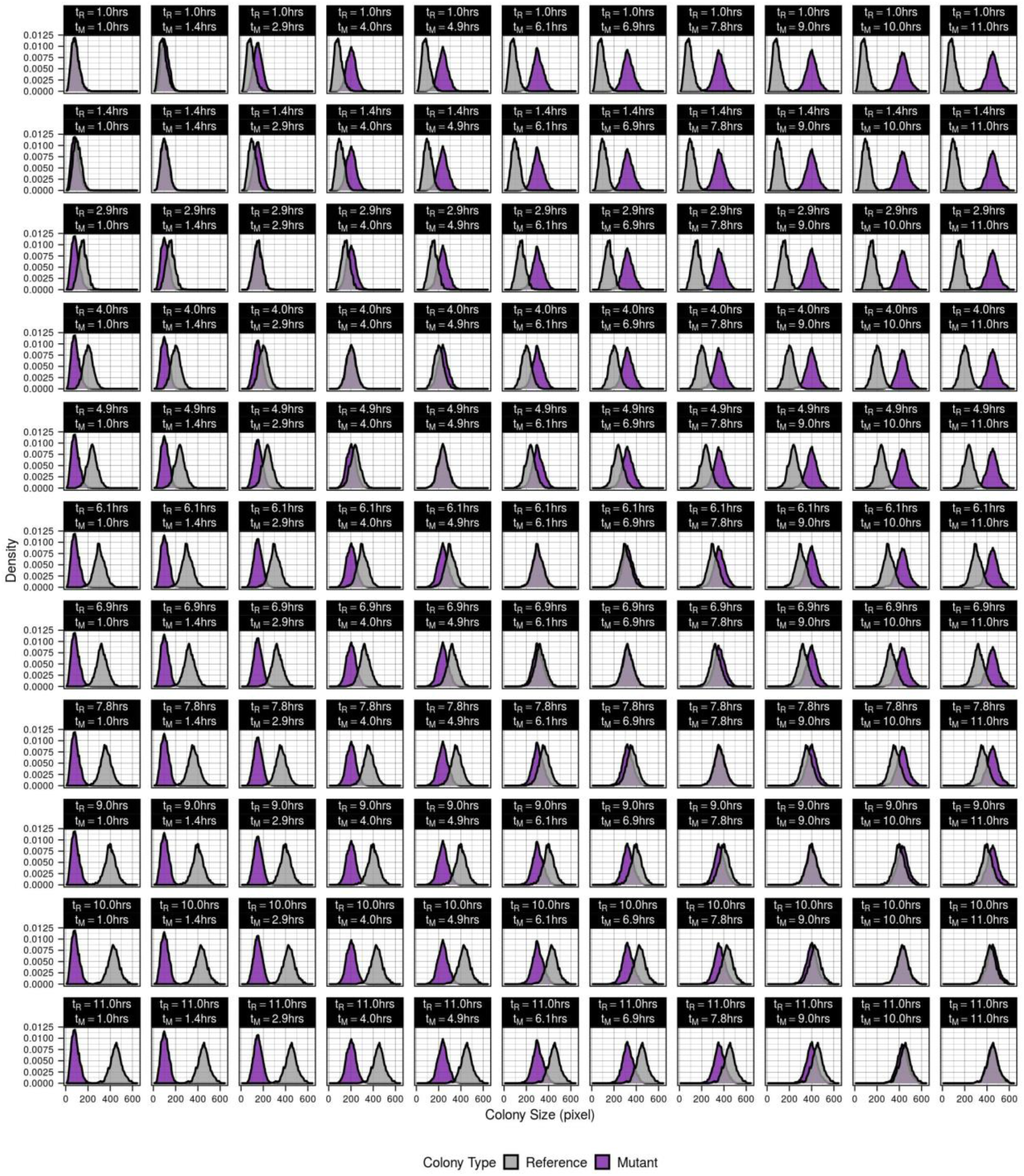
Colony size distributions of all virtual plates with a bimodal distribution. Composite density plot with each panel representing an individual plate. In each panel, the colony sizes of the reference population are represented by the grey density plot and that of the mutant by the purple density plot. The reference colony time point (t_R_) increases as we go from the top to bottom, and the mutant colony time point (t_M_) increases from left to right. The diagonal represents the real plate data where both the references and mutants are at the same time point – condition negative dataset. Everything to the left of the diagonal is condition positive deleterious data, and everything to the left is condition positive beneficial. **Figure 2C** is the fitness effect matrix of this composite plot.

**Figure S5.**
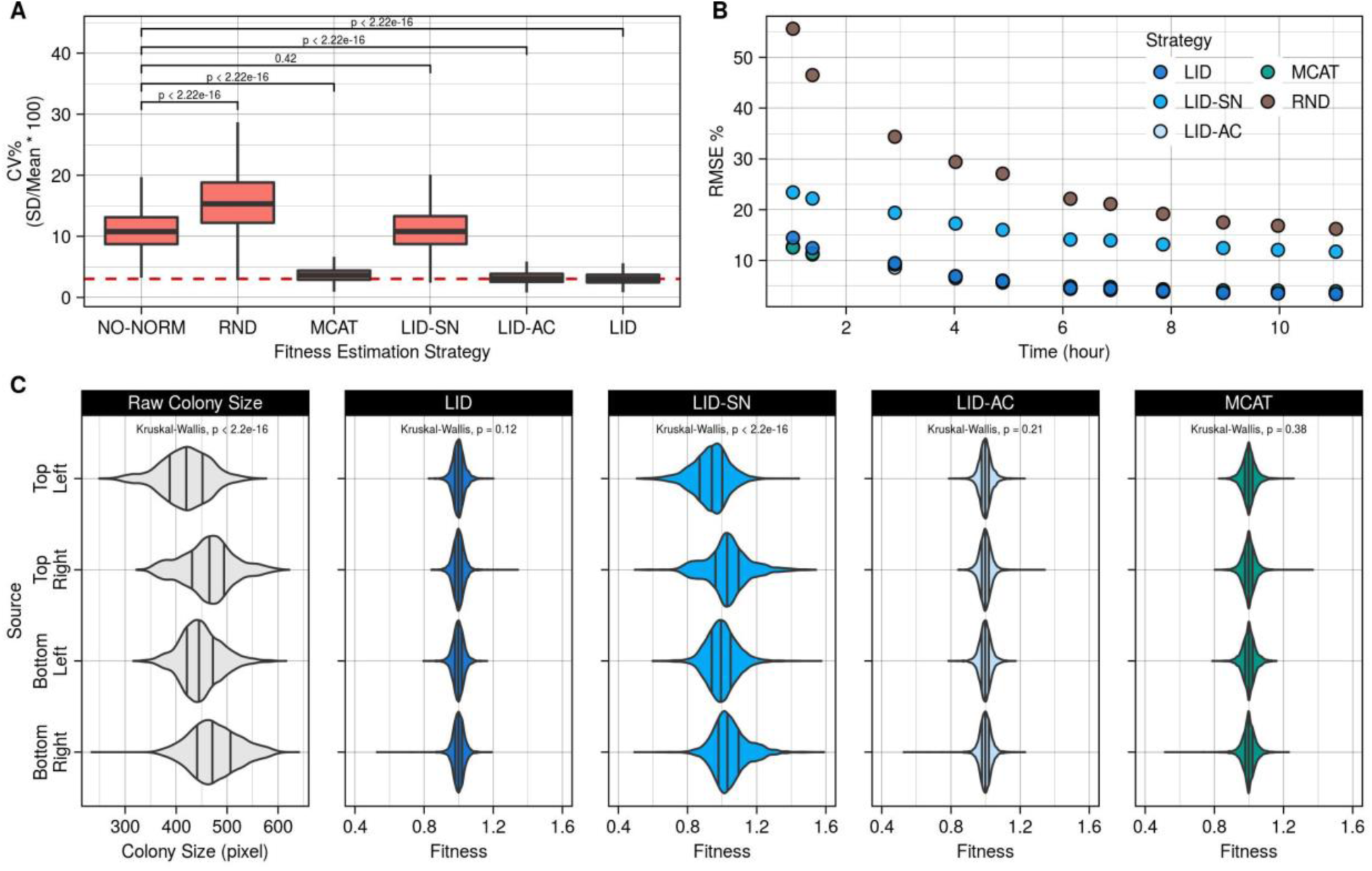
Accuracy of background colony sizes and source normalization. **A**. Coefficient of variance percentage (CV%) as a measure of spatial bias. The box plots show the CV% results for the raw colony size data (NO-NORM) and fitness estimated using a random colony size predictor (RND), Matlab Colony Analyzer Toolkit (MCAT)^14^, LI Detector without source normalization (LID-SN), LI Detector without local artifact correction (LID-AC) and LI Detector (LID). Wilcoxon rank-sum test was used to compare results between NO-NORM and the rest. **B**. Root mean square error (RMSE) of the background colony sizes compared to the observed colony size as a percentage of mean observed colony size per time point. Colors represent different strategies for fitness estimation. **C**. Source-plate-wise violin plot of raw colony sizes and LID, LID-SN, LID-AC, and MCAT^14^ normalized fitness at saturation (time = 11.0 hours). Solid black vertical lines indicate lower, middle, and upper quartile. The source-wise distributions are compared using a non-parametric (Kruskal-Wallis) test.

**Figure S6.**
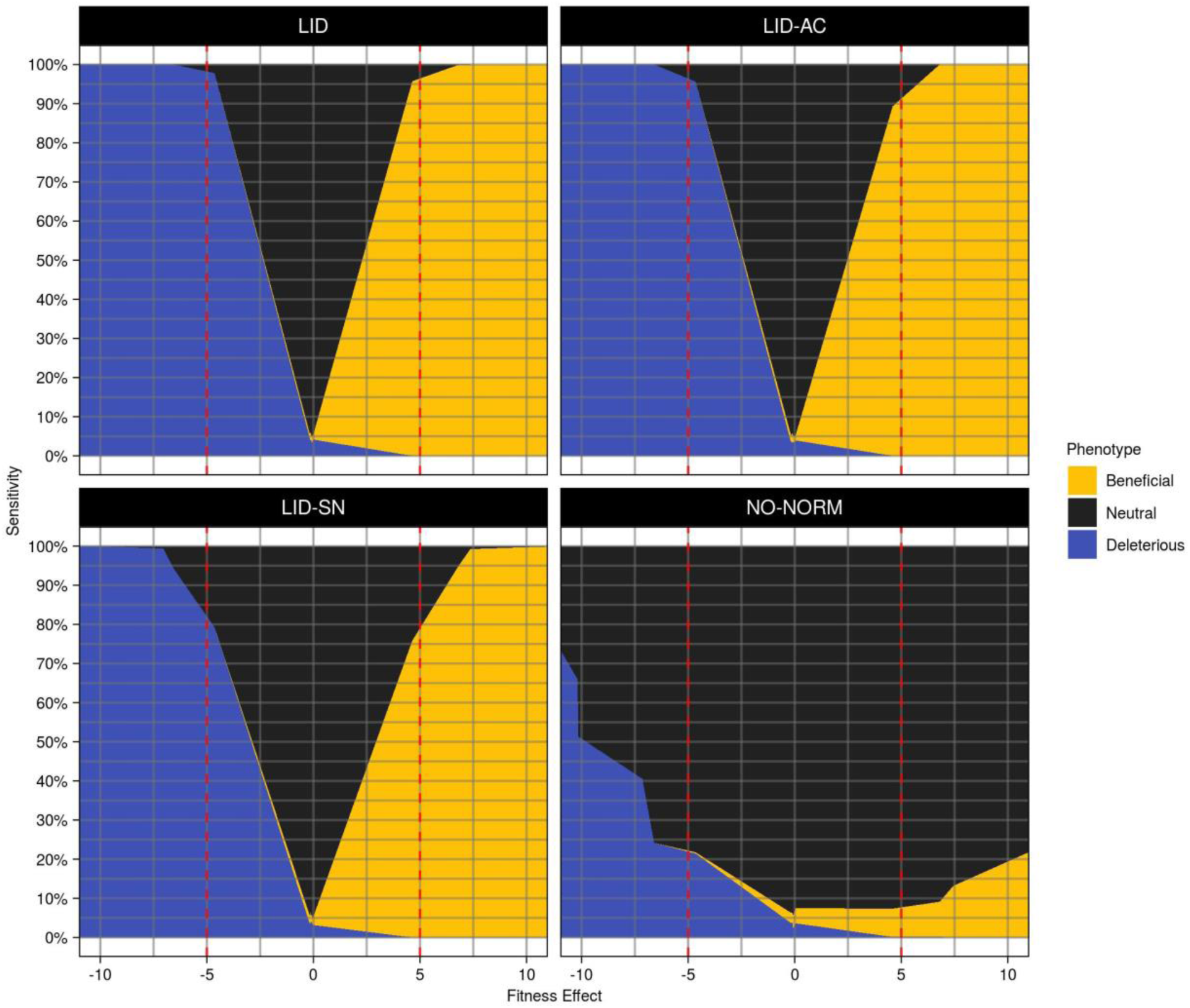
Sensitivity and different strategies for fitness estimation. Phenotype classification results from the virtual plates with bimodal distributions at a false positive rate of 5% are arranged according to increasing fitness effects. The dotted red line indicates a 5% fitness effect. Individual panels represent distinct strategies for fitness estimation (**see Materials and Methods**) - LI Detector (LID, top left), LI Detector without local artifact correction (LID-AC, top right), LI Detector without source normalization (LID-SN, bottom left), and no normalization (NO-NORM, bottom right).

**Figure S7.**
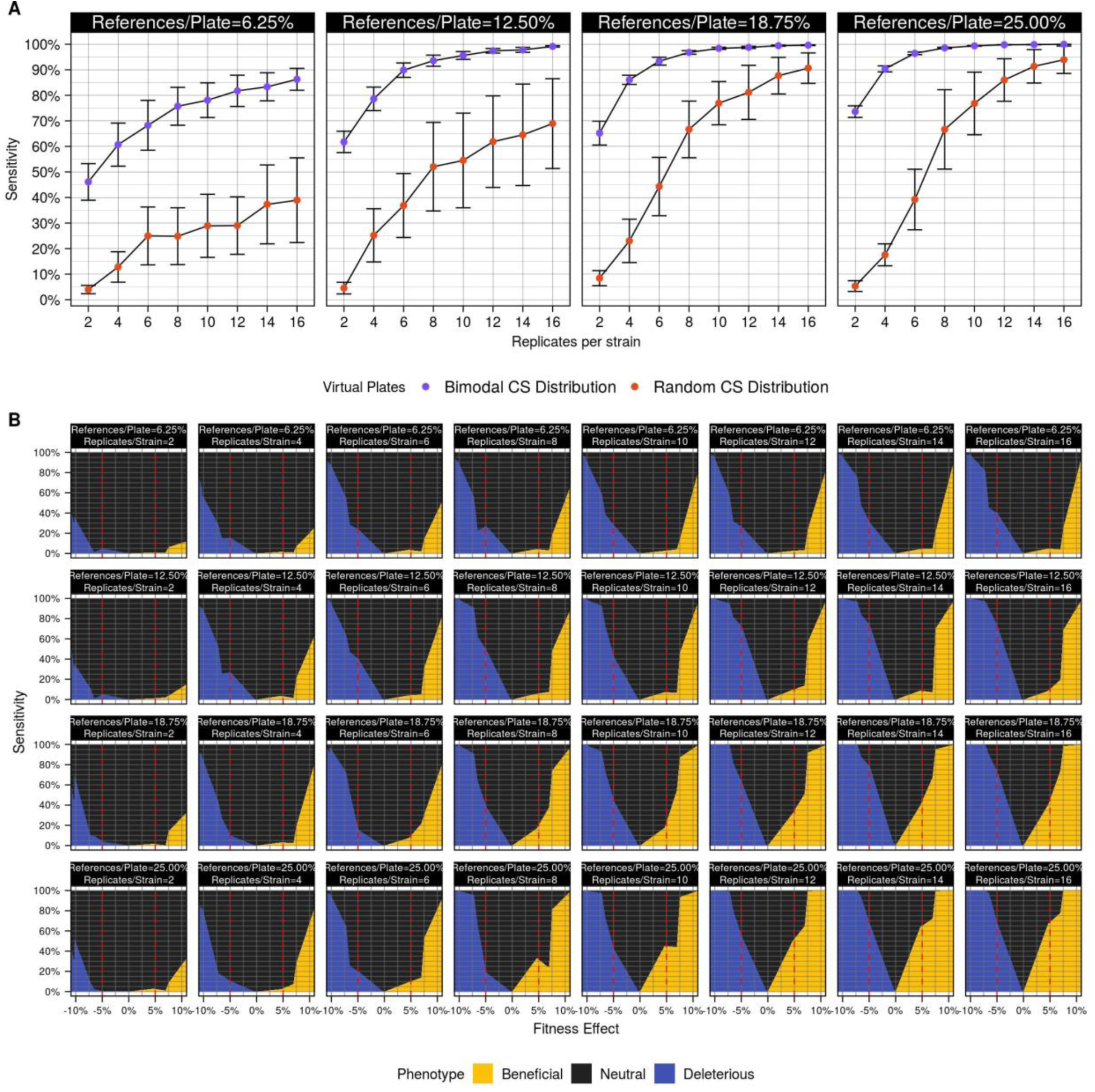
Sensitivity is directly related to the number of references and replicates. **A**. Sensitivity for observing 7% fitness effects, as a function of the varying proportion of references per plate (individual panels) and the number of replicates per strain (x-axis) was estimated for virtual plates with bimodal (purple) and random (orange) colony size distribution. Error bars represent a single standard deviation. **B**. Phenotype classification results from the virtual plates with random colony size distributions at a false positive rate of 5% are arranged according to increasing fitness effects. The dotted red line indicates a 5% fitness effect. Panels are arranged according to the increasing proportion of references per plate (top to bottom) and replicates per strain (left to right).

**Figure S8.**
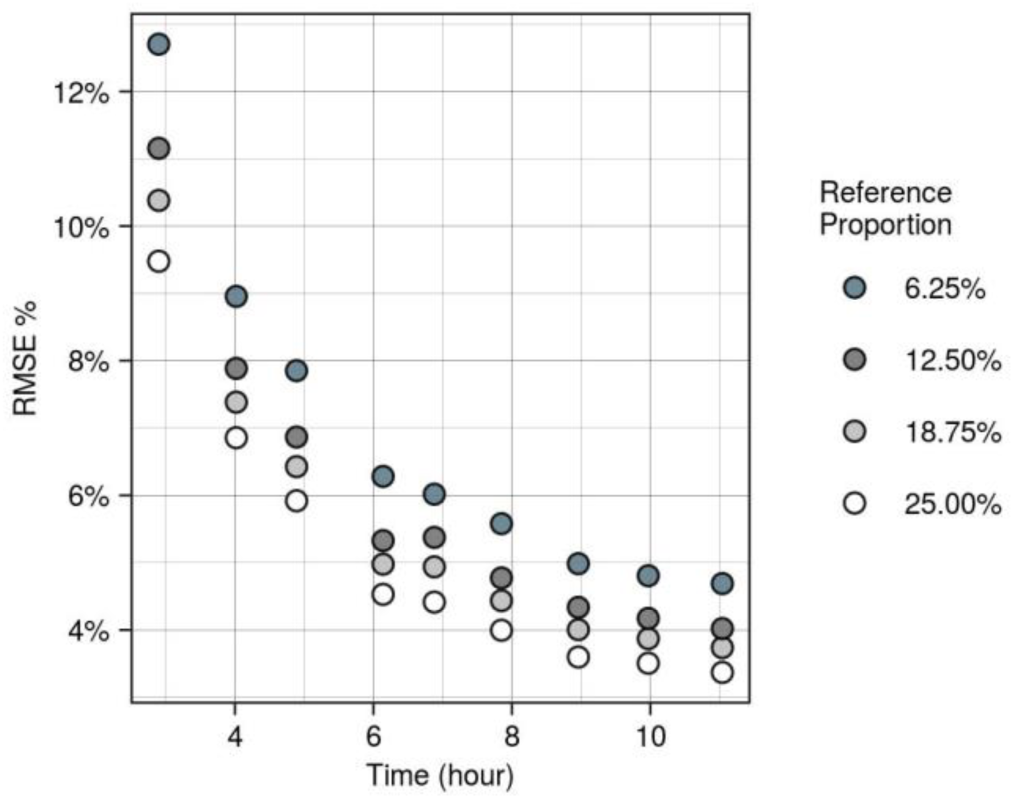
Effect on RMSE due to the proportion of references. The root mean square error percentage (RMSE%) for different proportions of references (colors) per time point was used as a measure of the accuracy of LI Detector’s reference-based background colony size. RMSE% decreases as colonies reach saturation and with an increasing proportion of references.

**Figure S9.**
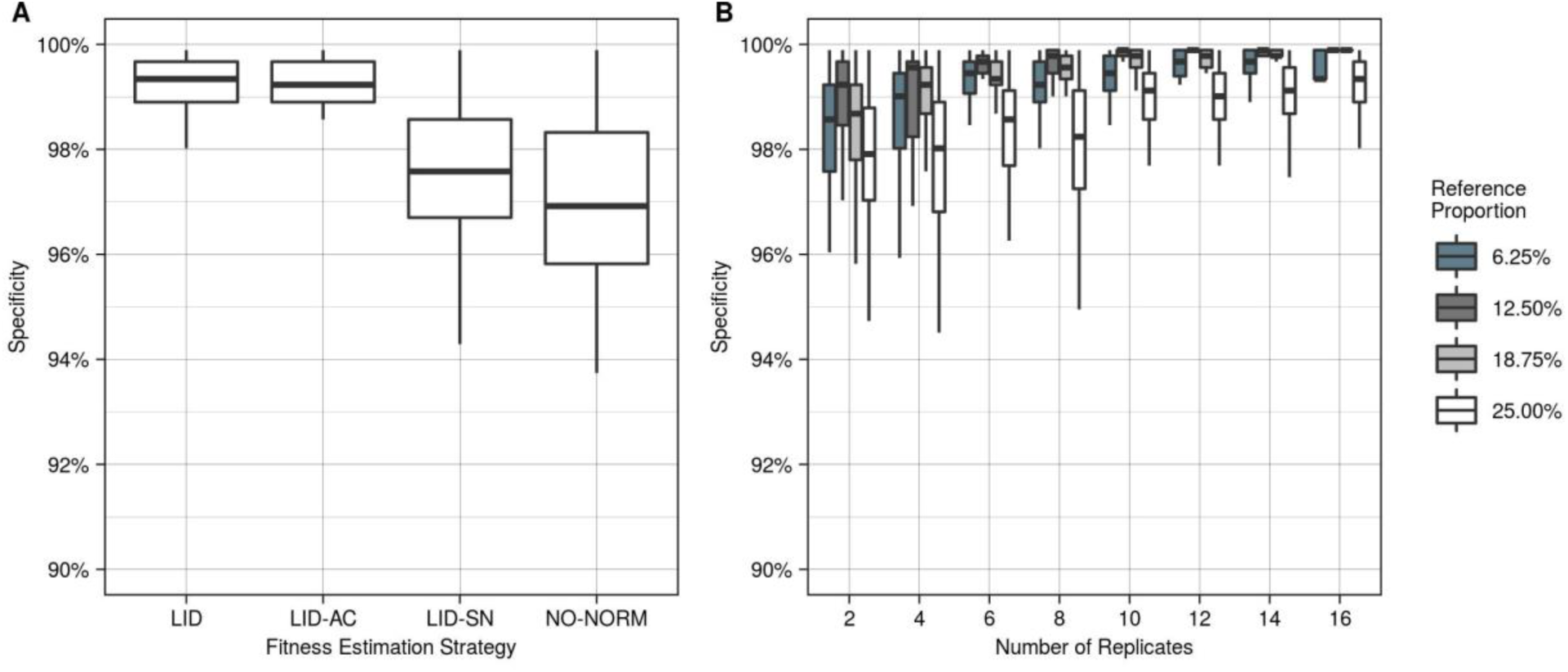
Specificity of different ways of using the LI Detector. Condition negative dataset used to measure the change in specificity with **A**. different strategies for fitness estimation (**see Materials and Methods**) consisting of LI Detector (LID), LI Detector without local artifact correction (LID-AC), LI Detector without source normalization (LID-SN), and no normalization (NO-NORM), where boxplots show pooled specificity results from all time points; and **B**. number of replicates for mutant and proportion of references per plate.

**Figure S10.**
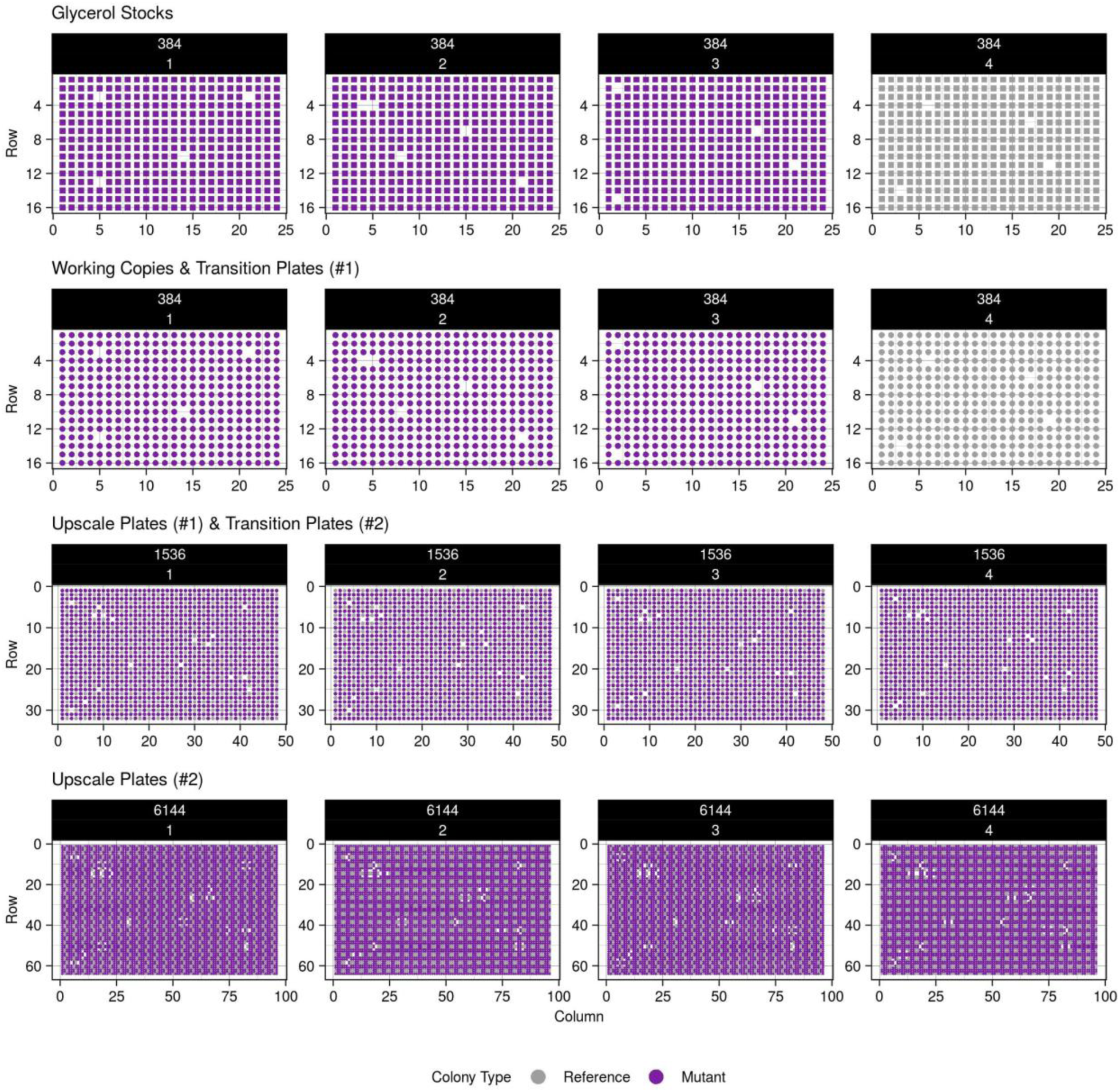
Visual representation of the plate maps. Spatial layout of colonies across the plates of the experiment. Each row represents a different stage/density of the validation experiment. These maps are made using information like plate number, column number, row number, strain identifier number, mutant name, and unique numeric identifier for the position on the plate stored as the pos2coor and pos2orf_name tables. For simplicity, the mock mutants are given a binary color classification of either reference or mutant.

**Figure S11.**
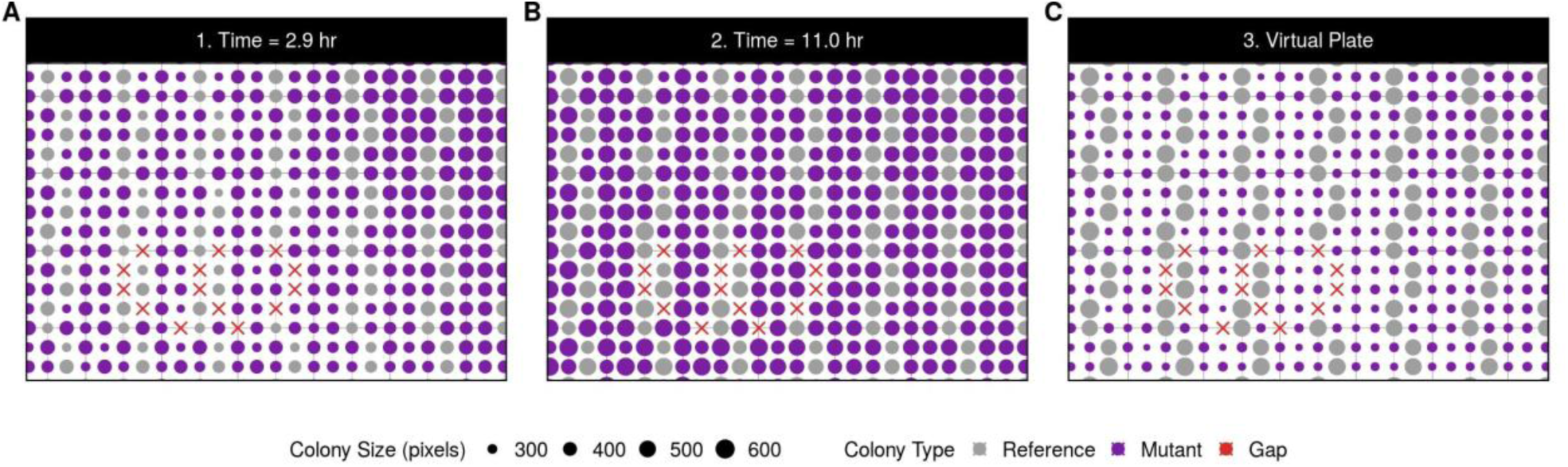
Making of a virtual plate with a bimodal distribution. The time-lapse images of the final screen plates can be used to create virtual plates where reference and mutant colony sizes come from different time points. The above three panels are a zoom-in version of the same region of a 6144-density plate. Colony type is represented by colors, and colony size by point size. **A**. Shows the colony layout and colony size estimations at 2.9 hours, **B**. shows the same region at 11.0 hours, and **C**. is an example of the virtual plate that can be created when the reference colony size data is taken from **B**. and mutant colony sizes from **A**. This plate still maintains the overall topological relationships of the colonies. In this example, all mutant colonies on the virtual plate are true positive deleterious by design.

